# Rapid, Efficient *Candida albicans* Lysis Comparison Using the Cellometer X2 Fluorescent Cell Viability Counter

**DOI:** 10.1101/2025.08.12.669735

**Authors:** Peishan Xie, Leo L. Chan, Mackenzie Pierce, Bo Lin, Yanhong Tong, Macy W. Veling

**Affiliations:** Revvity Health Sciences Inc., Hopkinton, MA; Revvity Life Sciences Inc., Lawrence, MA

**Author notes:** **Author Contributions** M.W.V. and Y.T. conceived the study. P.X. and M.W.V. performed the key experiments, carried out the analysis and drafted the paper. L.C., M.P. and B.L. assisted in cell counter setup and image analysis. All authors reviewed the paper.

**Keywords:** Cell Lysis, Cellometer X2, Fluorescent microscopy, qPCR, *Candida albicans*

## Abstract

Cell lysis is essential for extracting intake genetic material, forming the basis for diagnostic tests, and genetic studies. Commonly used lysis methods include thermal lysis, mechanical force, chemicals, biologicals, and sonication. Determining effective lysis methods for specific cell types is crucial for conducting further research. This study evaluates the lysis efficiency of *Candida albicans* using the Cellometer™ X2 fluorescent cell viability counter, employing various lysis methods: thermal, chemical, and enzymatic. Our results indicate that high-pH buffers combined with heat treatment enhance lysis efficiency, while SDS alone or with Proteinase K is insufficient for lysis at room temperature. In contrast, Zymolyase effectively lyses *C. albicans* at room temperature in 150 minutes, making the automation of room temperature lysis and nucleic acid purification feasible for *C. albicans*. Overall, this study highlights the Cellometer X2’s capability for rapid and direct evaluation of cell lysis efficiency.

## Introduction

Cell lysis is a crucial step in nucleic acid detection as it effectively breaks down cellular membranes to release DNA or RNA inside the cells. This process ensures that the genetic material is accessible for subsequent analysis, enabling accurate detection and quantification of nucleic acids. Without efficient lysis, incomplete or biased results, such as false negative detection, could occur. Hence, cell lysis lays the foundation for reliable diagnostic tests, research applications, and genetic studies. A fit-for-purpose lysis method with easy visualization/detection is essential for laboratories conducting any research activities involving nucleic acid and protein detection or cell membrane integrity measurement. Attractive lysis methods that have been broadly used include thermal lysis, mechanical force, chemicals, biologicals, or sonication (a combination of chemical and physical methods) ^[1]^.

Existing efficiency measurement methods can be categorized into two major characteristics: phenotype or genotype, each with its own strengths and weaknesses. Nucleic acid-based detection is a well-established genotypic method for lysis efficiency studies. However, it is sometimes not feasible for direct measurement due to the presence of inhibitors. These inhibitory molecules are often part of the sample preparation and nucleic acid extraction processes, interfering with the downstream nucleic acid detection ^[2]^. As a result, sample purification is often needed prior to detection, which could be complex and in turn, introduce process variation. Alternatively, in the phenotypic approach, direct lysis efficiency can be measured by image-based technology. Traditional bright-field microscopy is cost-effective but limited to only one staining dye, and it has relatively low resolution. Multi-color fluorescent microscopy, such as confocal, can provide more biophysiological information at cellular and sub-cellular levels with high resolution and imaging depth, while its price and complexity are not necessarily needed for a simple lysis efficiency study ^[3]^. Cellometer™ X2 fluorescent cell viability counter is a simple dual-fluorescent-based cell counting benchtop instrument, which is generally used for measuring cell stock concentration, viability, and cell size/diameter ^[4]^. It offers significant advantages over traditional bright-field microscopy, particularly in user-friendliness and efficiency. While professional multi-color fluorescence microscopes or confocal instruments are complex and require specialized training, the dual-fluorescent counter is intuitive and easy to operate. This makes it an ideal choice for direct cell lysis efficiency measurement, allowing researchers to quickly and accurately assess cell viability and functionality without the need for extensive technical expertise.

Candidiasis is a fungal infection caused by *Candida* species, and it is one of the leading causes of nosocomial bloodstream infections in the United States. Each year, an estimated 25,000 cases of candidemia, the bloodstream infection with *Candida*, occur in the United States with a mortality rate of about 25% ^[5]^. Among *Candida* species, *Candida albicans* (*C. albicans*) is a prominent yeast model organism and the most common cause of severe candidemia. Its robust cell wall consists of serval components, such as glucan and chitin, which are resistant to many existing mammalian or bacteria cell lysis methods ^[6]^. This presents unique challenges for cell lysis, making it an important target for optimization strategies. While the “gold standard” diagnosis method of invasive candidiasis has been culture-based, this approach has notable disadvantages, such as long turnaround time, and the risk of invasive sampling technique poses to some patients ^[7]^. Other non-cultural methods include tests based on nucleic acids, proteins, and metabolites ^[8]^; all rely heavily on the successful lysis of *C. albicans* cells. By optimizing lysis methods tailored to combat the resilient cell wall structure of *C. albicans*, researchers can extract high-quality cellular components for further studies. This is key to researching and developing diagnostic tools, ultimately aiding in the effective treatment and management of candidiasis outbreaks in public health ^[8]^.

Here, we utilized the Cellometer™ X2 fluorescent cell viability counter, to support *C. albicans* direct lysis efficiency studies. Lysis methods from thermal lysis, chemical-based buffers, and enzymatic reagents of choice are demonstrated in the study. Cell concentration and viability results suggested that the combined effect of high-pH buffers and heat treatment enhances lysis efficiency, while Zymolyase can effectively lyse *C. albicans* at room temperature (RT) in 150 minutes.

## Materials and Methods

### *C. albicans* culturing

*C. albicans* frozen stock received from BEI (Cat. #NR-29445) was seeded on a Candida BCG agar plate (Edge Biologicals) for one day at 27°C. After the formation of visible colonies, they were collected and resuspended in 700 µL of 1X PBS buffer (Thermofisher, Cat. # 10010023). The starting culture stock concentration was defined as 100X, approximately 6 × 10^8^ cells/mL.

### Cell staining and quantification

At each testing time point of all treatments, the treated sample was mixed with yeast acridine orange/propidium iodide (AO/PI) dye (Revvity, Cat. # CSK-0102-2mL) at a 1:1 ratio. 20 µL of the dye-mixed sample was then loaded into a disposable counting chamber (Revvity, Cat. # CHT4-SD100-002) and subsequently inserted into the Cellometer X2 instrument (Revvity, Cat. # CMT-X2-S150). The cells were allowed to settle inside the chamber for 5 minutes of incubation prior to image acquisition and analysis. Cellometer X2 employs bright-field and dual-fluorescent imaging modes to automatically determine the stock concentration, viability, and fluorescent intensity of target cells.

### pH-based chemical and thermal combined lysis

The 100X *C. albicans* culture in 1X PBS buffer was diluted to 10X by Buffer A (pH 8.0, 10mM Tris-HCl, 1 mM EDTA), Buffer B (pH 9.5, Revvity, Cat. # CSK-0102-2mL) and Buffer C (pH 12, 30mM Tris, 20mM KOH). Heat treatment was applied at 95°C for 5, 10, or 20 minutes (min) to each sample. Controls at 0 min without heat treatment for each lysis buffer type were included for comparison. After incubation, the cell staining, and quantification procedure described above were followed.

In the qPCR experiment, a serial dilution of the 100X culture in 1X PBS buffer was performed to reach final concentrations at 20, 2, 0.2, and 0.02X. 50 µL of each sample was incubated either at 95°C or RT for 5 mins. 5 µL of each sample was loaded to a final 20 µL PCR reaction (Promega, Cat. # M7405). A set of primers/probe specific to a *C. albican*s target gene, which is present as a single copy in the genome, was applied in the PCR reaction ^[9]^. Three replicates per testing condition were included. QuantStudio™ Dx 96 (ThermoFisher) was used with real-time cycles run at 94°C/2 mins, 40 cycles of 94°C/10 sec, 62°C/15s, and 65°C/45s for signal collection.

### Detergent-based chemical lysis

The 100X *C. albicans* culture in 1X PBS buffer was first diluted to 50X by 1X PBS buffer, followed by a second round of dilution to 25X by buffer (10mM Tris-HCl, 1 mM EDTA, pH 8∼8.5), buffer with 1% SDS, or buffer with 1% SDS and Proteinase K (18 mg/mL). After a second round of dilution, 200 µL of the sample contains approximately 1.5 × 10^7^ cells. These samples were then incubated at RT throughout the whole process. At each time point (0, 15, 30, 60, and 90 min), the cell staining, and quantification procedure described above were followed.

### Enzymatic-based lysis

The 100X *C. albicans* culture in 1X PBS buffer was first diluted to 50X by 1X PBS buffer, followed by a second round of dilution to 25X by TE buffer (pH 8.0, 10mM Tris-HCl, 1 mM EDTA) or TE buffer with Zymolyase (Zymo Research, Cat. # E1004). After a second round of dilution, 50 µL of the sample contains approximately 4.3 × 10^7^ cells with or without 5U Zymolyase. These samples were then incubated at RT throughout the whole process. At each time point (0, 30, 60, 90, and 150 min), the cell staining, and quantification procedure described above were followed.

In lysis efficiency comparison between Zymolyase and Proteinase K, a 3-fold serial dilution of the 100X fresh culture by liquid amies (BD, Cat. # 220245) was performed to reach final concentrations at 1, 1/3, 1/9, and 1/27X. The 300 µL of lysis buffer and 10 µL of Proteinase K from the chemagic™ Pathogen NA Kit H96 (Revvity, Cat. # CMG-1033-G) were used as “ProK (+)” solution according to kit manufacturer’s guideline. The same amount of lysis buffer from the chemagic kit with Zymolyase (1U/rxn) spike-in was set up as “Zymolyase (+)” solution. 310 µL of each prediluted sample was spiked into “ProK (+)” and “Zymolyase (+)” solutions, and negative controls, *C. albicans* (-) ProK (+) and *C. albicans* (-) Zymolyase (+), were also prepared. Samples were incubated at RT for 30 minutes, followed by an automated extraction procedure on the chemagic^™^ 360 instrument (Revvity). Six replicates per testing condition were included. 10 µL of the extracted DNA was loaded to set up a final 15 µL PCR reaction. The Reagent A (PCR buffer) and Reagent C (enzyme mix) from the *Candida auris* Detection Real-time PCR Reagents (Revvity, Cat. # DXMDX-RGT-1001) with *C. albican*s primers/probe set were used to set up the reaction. CFX 96 (Bio-Rad) was used with real-time cycles run at 37°C/2 mins, 94°C/10 mins, 42 cycles of 94°C/10 sec, 62°C/15s, and 65°C/45s for signal collection signal record).

## Results

### Combined high pH and heat enhance *C. albicans* lysis

Three detergent-free buffers, Buffers A, B, and C, ranging from mild to high pH (pH 8.0, 9.5, or 12), were used in combination with heat treatment (HT) at 95°C for 0, 5, 10, or 20 mins for *C. albicans* lysis efficiency comparison. The viability result, measured by the Cellometer X2 software, showed that the proportion of propidium iodide (PI)-positive *C. albicans* cells (in red: dead cell marker) increased from 8.37% to nearly 50% in the mild pH buffer (Buffer A) as the duration of heat treatment extended from 0 to 10 mins (Figure 1-2). Interestingly, the PI-positive cells (49.57% - 49.87%) were also acridine orange (AO)-positive (in green: live cell marker, 50% - 50.43%) in a 1:1 ratio between the 5- and 10-min heat-treat conditions. This indicated that the entire *C. albicans* population was close to 100% damaged, yet they were not completely lysed or destroyed. In contrast, the viability in both the medium and high pH buffers (Buffer B and C), with a 5-min heat treatment, showed a 0% AO marker, indicating 100% cell death. In fact, the PI marker was almost lost in Buffer C even at 5-min heat-treat condition when compared to Buffer B, suggesting higher pH further enhances cell destruction to completion. An additional control in Buffer C at 150 mins without heat treatment confirmed that only approximately 20% viability was reduced from 0 min (data not shown), suggesting that a buffer with high pH property alone is insufficient to destroy *C. albicans* structure, even after 3.5 hours of incubation.

**Figure 1.**
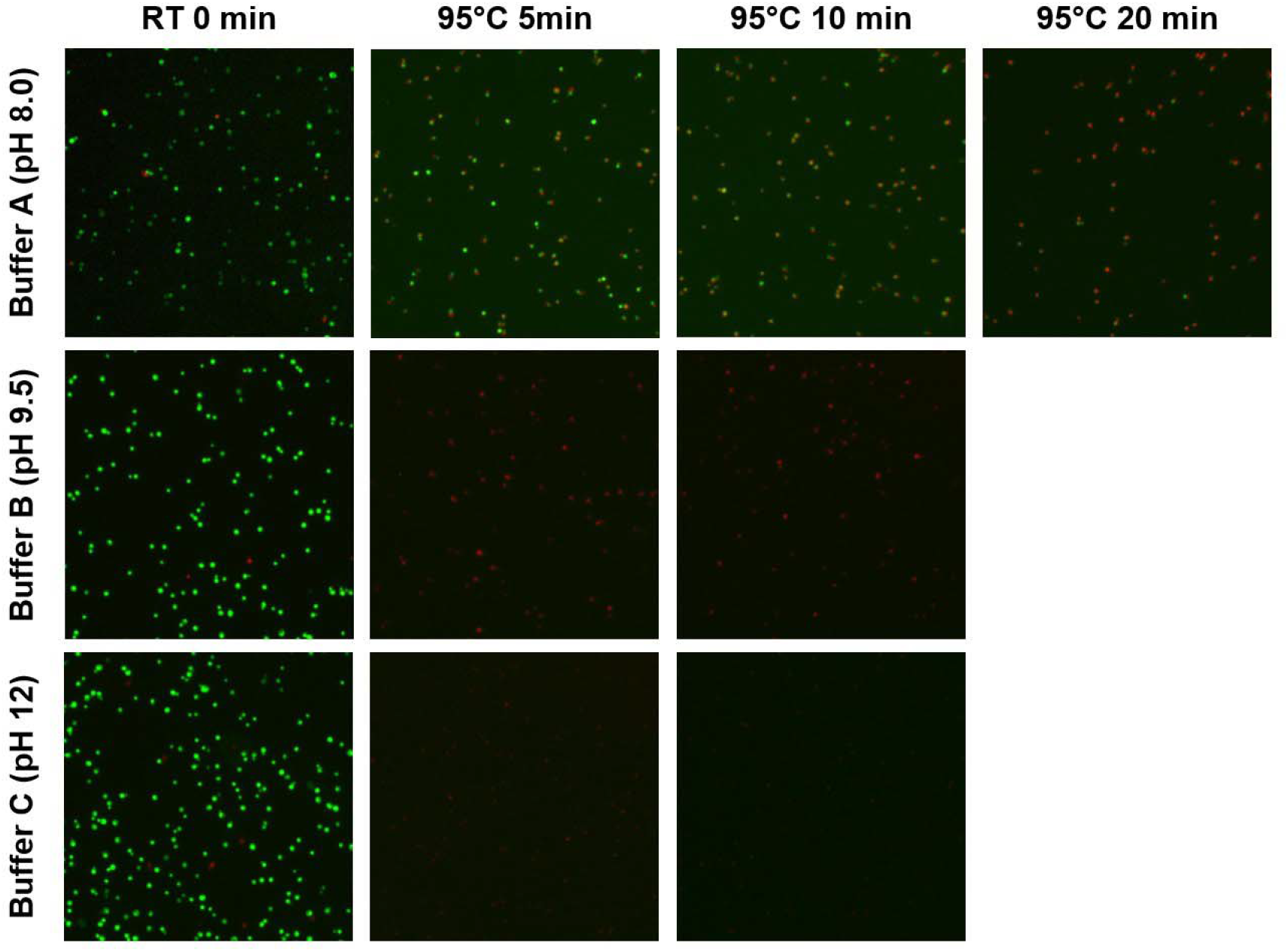
*C. albicans* AO/PI staining in buffers with different pH at various heat-treat conditions. Green: AO, live cell marker. Red: PI, dead cell marker. Yellow: merge of AO/PI, damaged cell marker with cell structure retained. RT: room temperature. Scale bar not shown.

**Figure 2.**
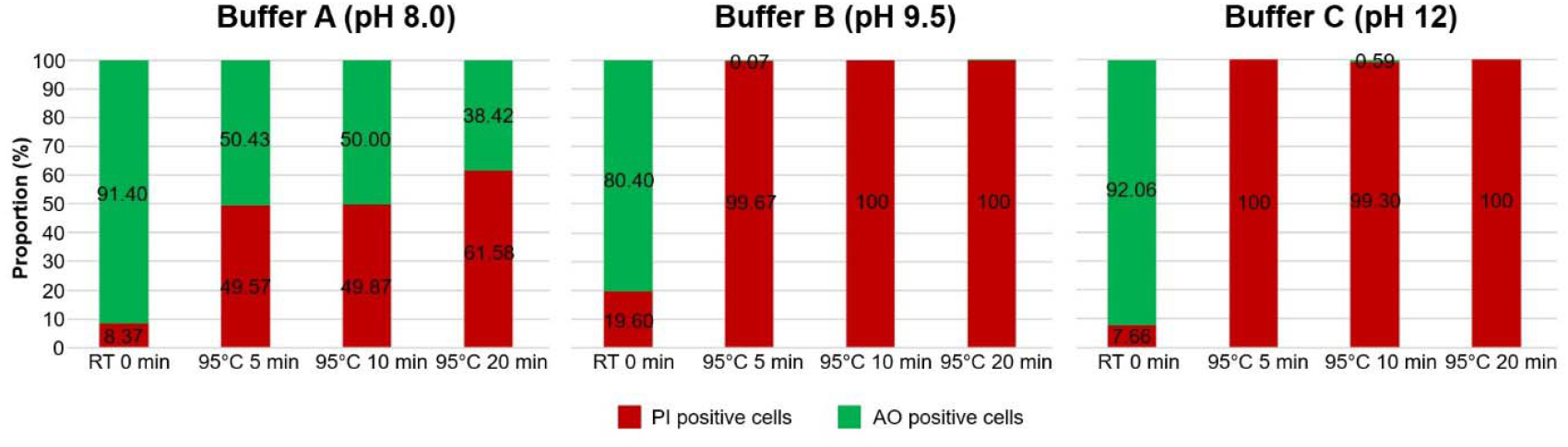
Proportion (%) of AO- or PI-positive cells in different lysis buffers at various treatment time points.

To confirm the phenotypic observation that a combination of higher pH and HT enhances lysis efficiency more effectively than either one, a genotypic confirmation using qPCR was applied. Given the same absolute input condition, qPCR results consistently showed that the higher pH buffer produces a smaller Ct value than the milder pH buffer, in both RT and HT cases. When the *C. albicans* concentration was at 0.02X, the sample treated with Buffer A failed to amplify (Figure 3). This suggests greater lysis efficiency and increased DNA release, confirming the same observation as measured by phenotype.

**Figure 3.**
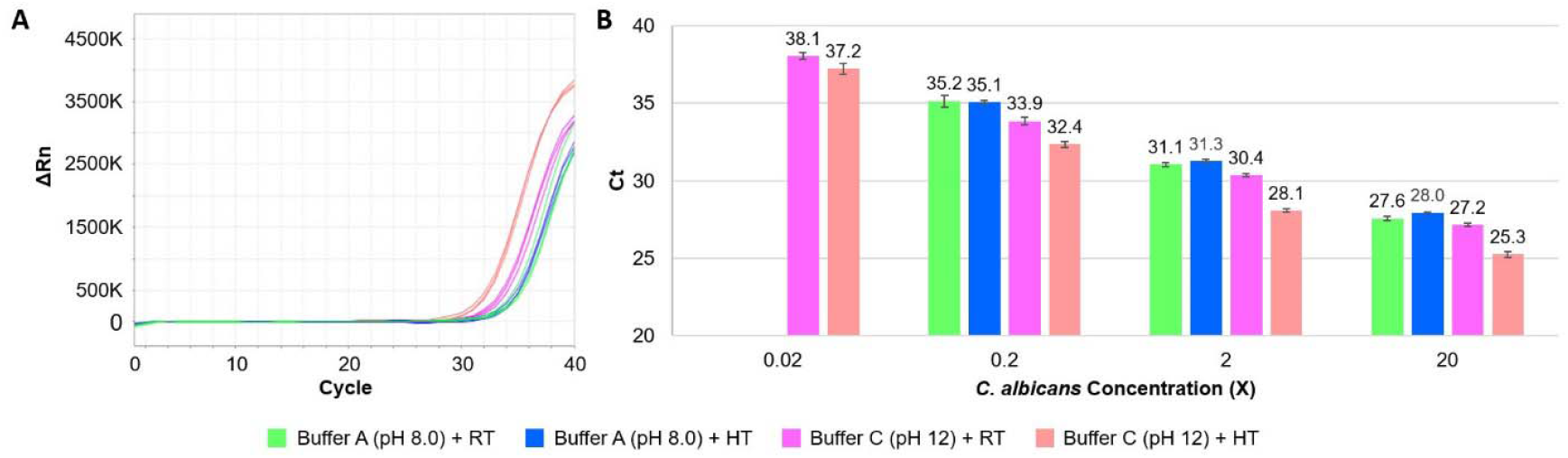
Realtime PCR quantification of *C. albicans* lysis efficiency by buffer A and C under RT or heat treatment (HT). **(A)** Amplification curve shown at 0.2X concentration as an example. **(B)** Mean Ct value of specific treatment cases at various concentrations (n=3). *C. albicans* Concentration (X) denoted sample treatment concentrations. *C. albicans* were not detected in Buffer A at the 0.02X concentration in either RT or HT. Error bars represent standard deviation.

Thermal disruption is commonly used in addition to either enzymes or mechanical beating and can significantly improve fungal DNA purification ^[10]^. Both phenotypic and genotypic methods in this study illustrated that the lysis efficiency of the higher pH condition is enhanced by high-temperature treatment.

### Detergent addition helps but is insufficient for *C. albicans* lysis

Although buffers with high pH under high-temperature treatment can effectively destroy the viability of *C. albicans* in 5 minutes, it would be more convenient if an RT solution is available to replace the high-temperature step. To explore an RT solution for thick yeast cell wall destruction, a mild basic buffer (pH 8-8.5) with strong detergent SDS was examined over a 90-min time course study. Proteinase K was also included in the study as it is commonly combined with SDS in many bacterial and mammalian cell lysis buffer recipes. The total cell (brightfield), AO-positive, and PI-positive cell numbers measured at each time point were normalized to pre-treatment sample concentration for comparison. In the Buffer-only condition, the cell numbers of total (brightfield), AO-positive, and PI-positive remained relatively similar from 0 to 90 mins (Figure 4 and 5A). In the Buffer with SDS, it led to a 4-fold increase in PI-positive cells compared to the control at 15-min of incubation. These PI-positive cells from 15-to 90-min are also nearly all AO-positive with their ratio close to 1:1 (Figure 5B). As the total cell numbers remained consistent over the study period, these observations suggested that the buffer with SDS at RT is sufficient to permeabilize the *C. albicans* cell wall at 15 mins but incapable of fully destroying the cell structure even after 90 mins incubation. Strikingly, adding Proteinase K to the SDS buffer didn’t seem to improve cell destruction at RT, at least not by phenotypic observation at a fixed time point (Figure 5). This study demonstrates that a buffer with SDS, in the presence or absence of Proteinase K, can effectively damage the cell wall of *C. albicans* at RT, but it is insufficient to fully lyse the cell structure even over a 90-min incubation. This study also demonstrates the potential of using the Cellometer X2 rapidly to evaluate lysis efficiency with detergent-based methods, particularly with strong detergents, which are typically not feasible for qPCR due to their inhibitory effects.

**Figure 4.**
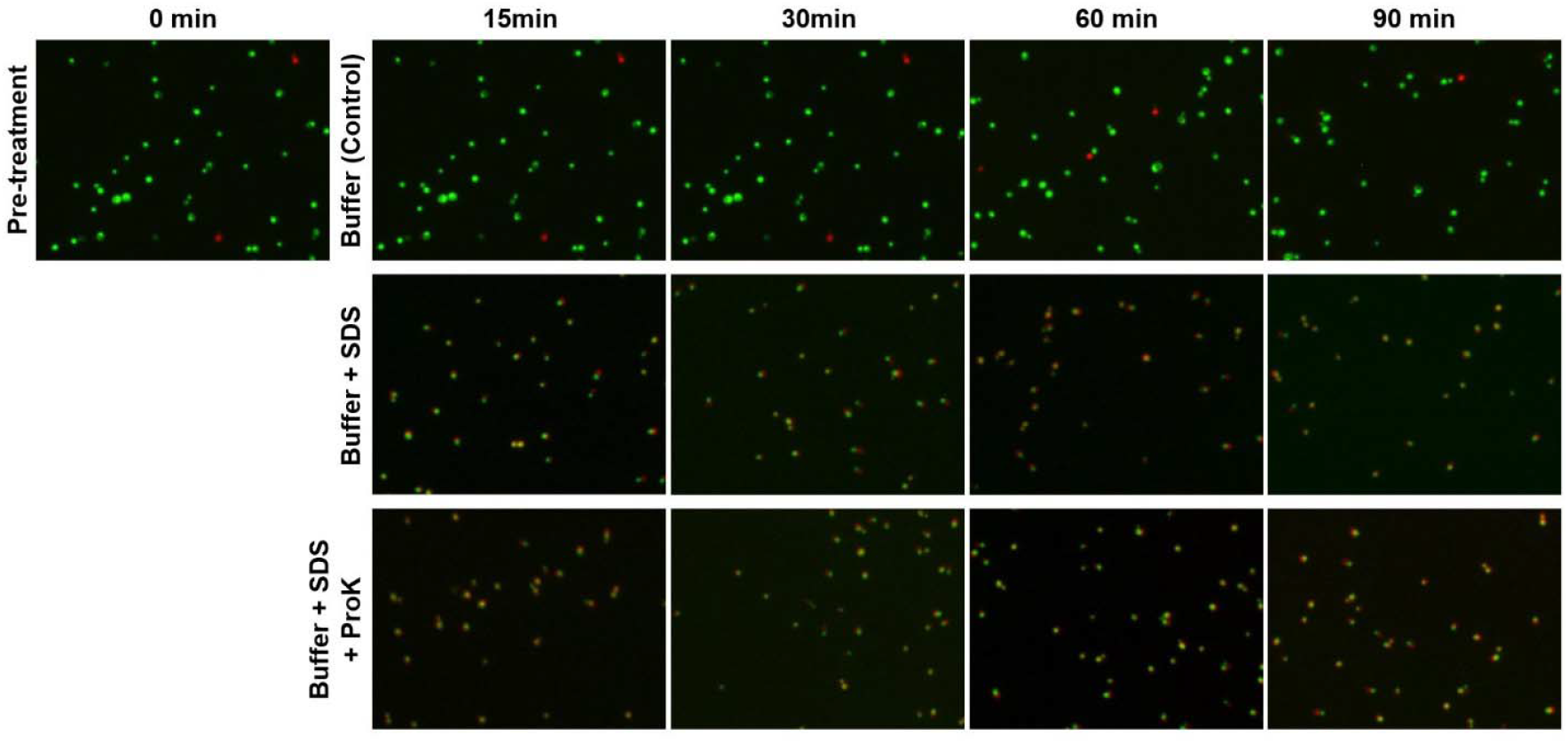
*C. albicans* AO/PI staining in control buffer, and SDS buffer with or without Proteinase K (ProK) over a 90 mins RT incubation. Green: AO, live cell marker. Red: PI, dead cell marker. Yellow: merge of AO/PI, damaged cell marker with cell structure retained. Scale bar not shown.

**Figure 5.**
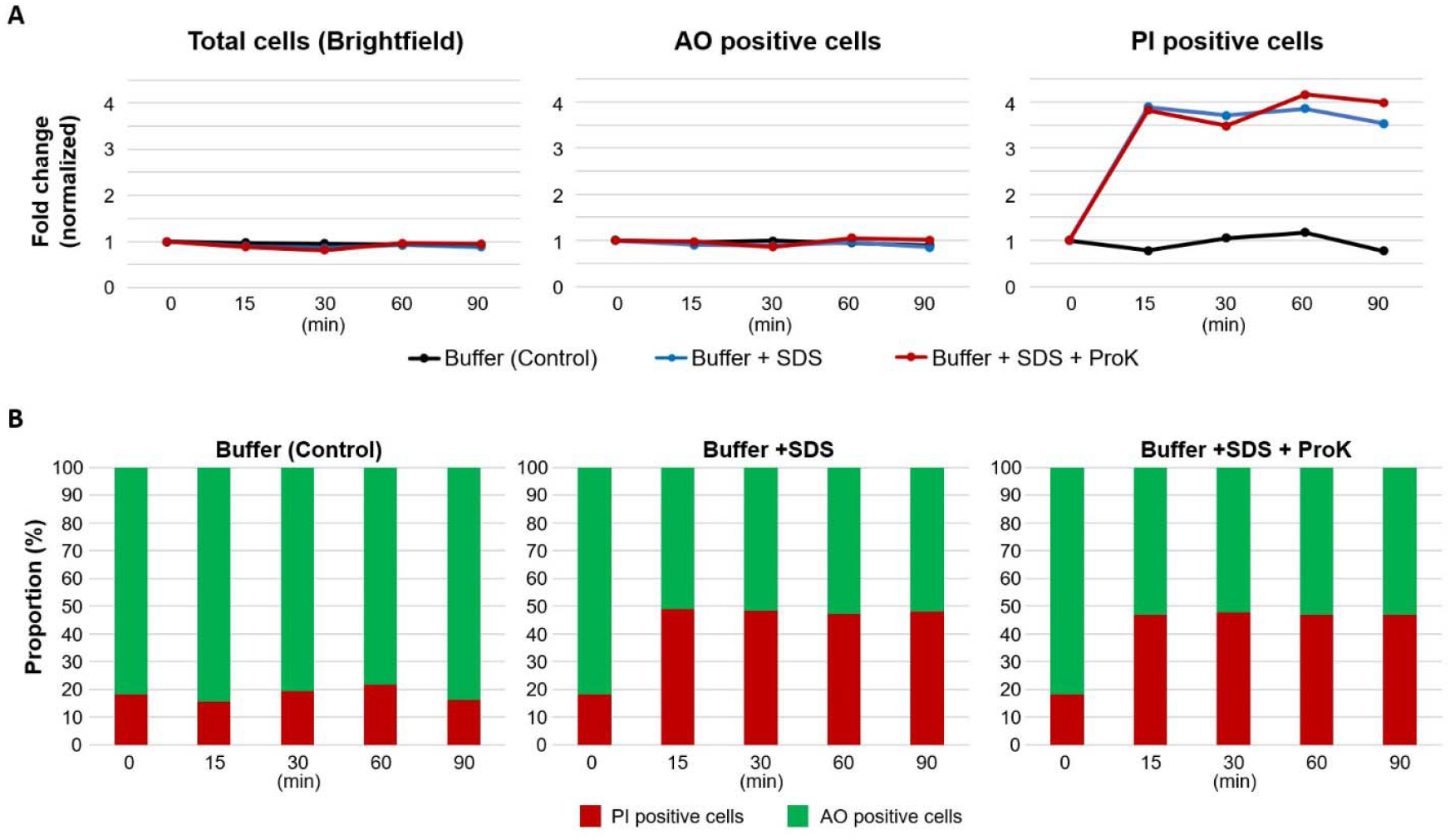
Lysis efficiency comparison of control buffer, SDS buffer, and SDS buffer with ProK over a 90 mins time course at RT. **(A)** Fold change of total, AO-positive, and PI-positive cells under each buffer type over time. **(B)** Proportion (%) of AO- or PI-positive cells in individual buffers over time.

### Yeast-specific enzymatic method offers a room-temperature solution for *C. albicans* lysis

Given that a buffer with strong detergent, in the presence or absence of Proteinase K, is insufficient to fully lyse *C. albicans* at RT, an alternative enzymatic method ^[11]^ using Lyticase, such as Zymolyase was evaluated over a 150-min time course study. Lyticase, which hydrolyzes glucose polymers at the β-1,3-glucan linkages in the yeast cell wall, has been suggested to improve lysis efficiency ^[12]^. Timelapse in conditions with or without the Zymolyase was performed for comparison (Figure 6A). The total cell (brightfield) and the AO-positive cell numbers measured at each time point were normalized to the same pre-treatment sample concentration for comparison. As a result, the proportion of total cells and AO-positive cells showed a similar degree of cell reduction pattern (Figure 6B). This enzymatic reaction was able to reduce total and AO-positive cell numbers down to half in 30 mins and further down to ∼5% in 150 mins. In contrast, the control showed that ≥ 70% of cells remained AO-positive even at 150 mins. These results suggested that Zymolyase was able to lyse *C. albicans* effectively at RT. Interestingly, the PI marker failed to provide quantitative information due to clusters of red signals, which likely were the leftover indigestible genomic material. Hence, it is not quantitative for total damaged or dead cell count.

**Figure 6.**
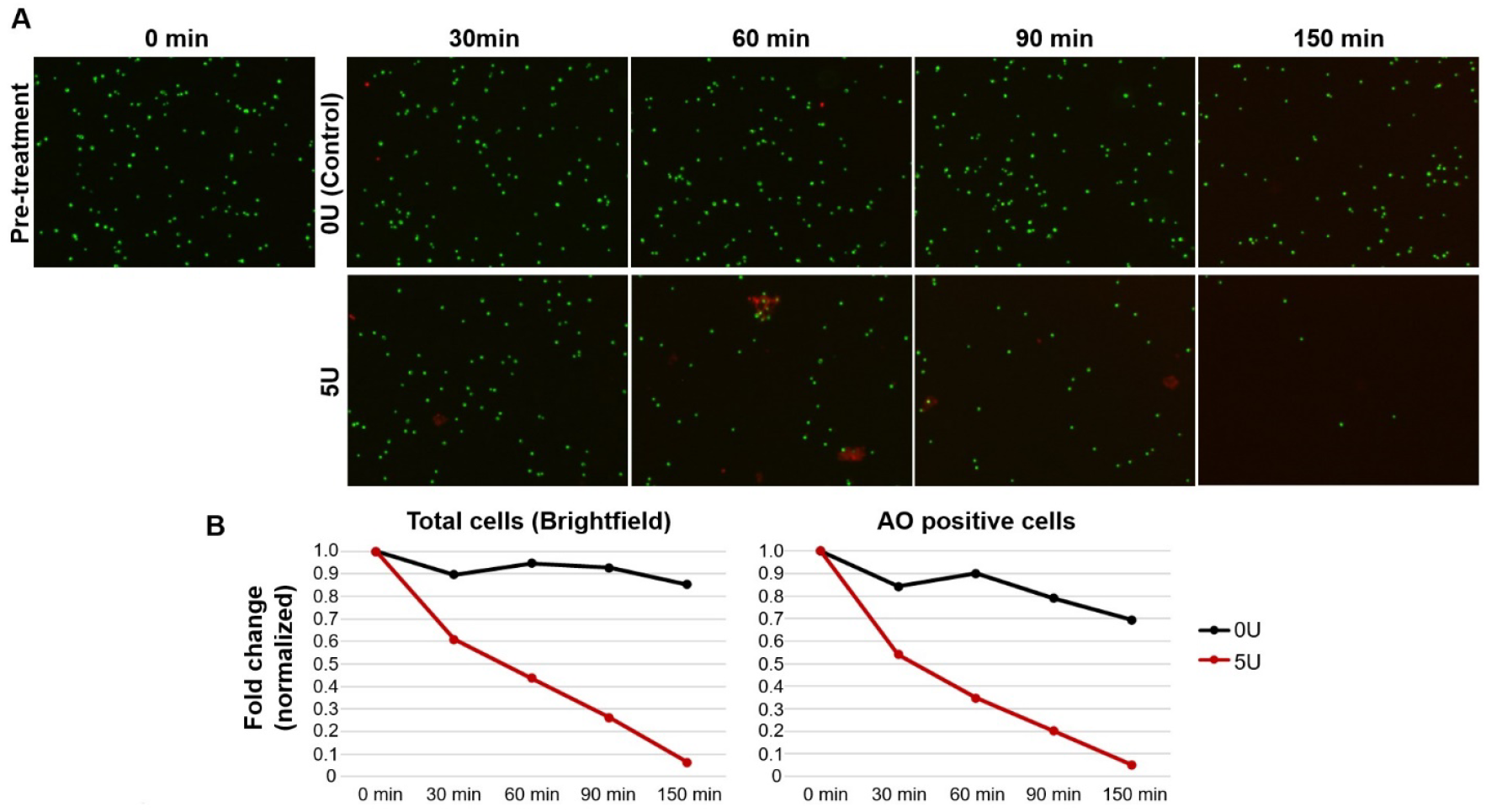
Lysis efficiency with or without Zymolyase over time at RT. **(A)** *C. albicans* AO/PI staining shown at different time points in both control (0U) and Zymolyase treatment (5U). Green: AO, live cell marker. Red: PI, dead cell marker. Scale bar not shown. **(B)** Fold change of total and AO-positive cells over time at each condition.

To confirm whether Zymolyase is more effective for lysing *C. albicans* than the Proteinase K Method provided by the chemagic™ Pathogen NA Kit H96 kit, different concentrations of *C. albicans* samples were prepared and incubated in lysis buffer (from chemagic™ Pathogen NA Kit H96 kit, see methods) containing either enzyme. Following this, all samples underwent extraction, and a qPCR study was performed. The results showed that the lowest detectable concentration of *C. albicans* is 9-fold more sensitive with Zymolyase than Proteinase K (Figure 7), confirming that Zymolyase indeed has better lysis efficiency than Proteinase K for treating *C. albicans*.

**Figure 7.**
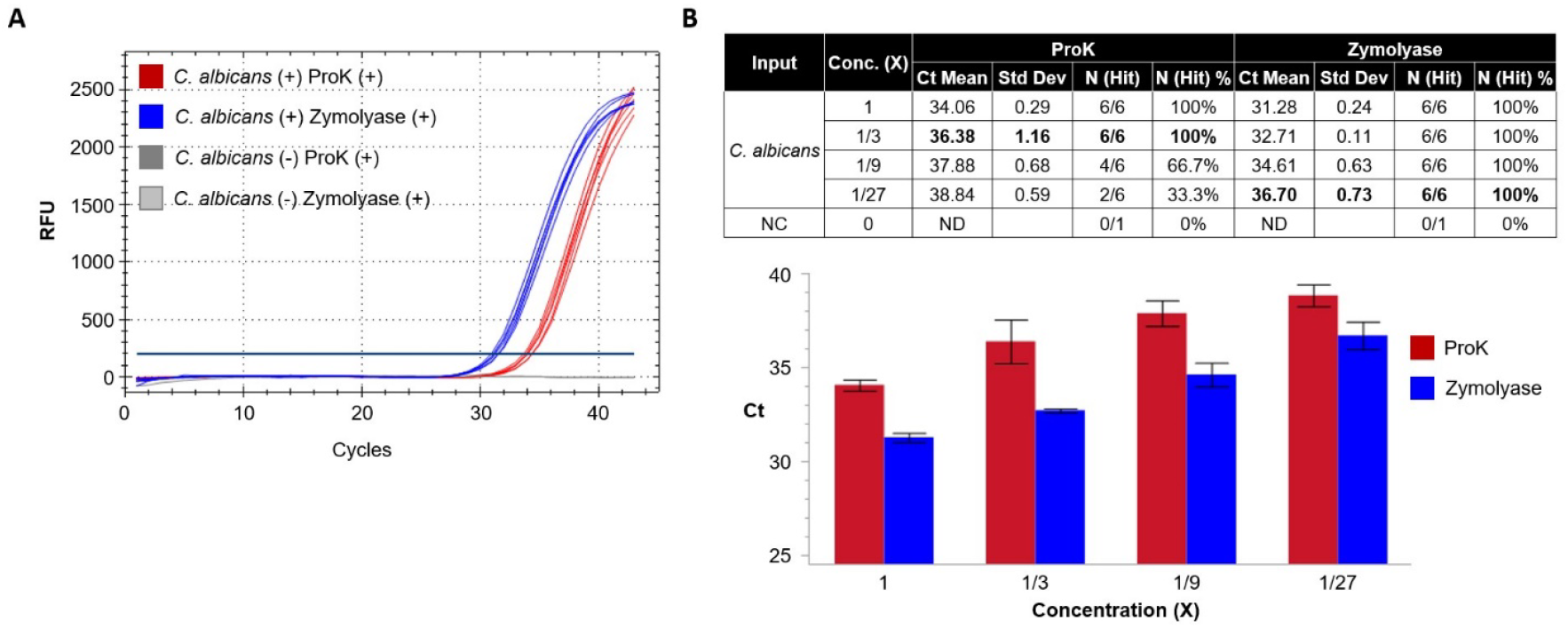
Lysis efficiency comparison of buffer with ProK or Zymolyase at RT by qPCR. **(A)** Example of qPCR amplification curve of buffer with ProK or Zymolyase at 1X concentration. **(B)** Table and bar chart showed the Ct mean value and detection rate of *C. albicans* at different concentrations between the two buffer conditions. NC: Negative control, including *C. albicans* (-) ProK (+) and *C. albicans* (-) Zymolyase (+). ND: Not detected. Error bars represent standard deviation.

Overall, both phenotypic and genotypic studies demonstrate that the enzymatic method using Zymolyase provides effective RT lysis on destroying *C. albicans* cells.

## Discussion

A successful RT solution for lysing *Candida* species is important, as standard lysis reagents designed for killing bacteria and mammalian cells may not be effective. Often, additional treatment steps are needed to enhance lysis efficiency. A simple, affordable platform like the Cellometer™ X2 fluorescent cell viability counter, together with the ViaStain Yeast Kit, can accelerate such studies by providing quick phenotypic observations in a timely manner. Using the Cellometer X2 together with an orthogonal qPCR assay, this study has demonstrated that high pH-based chemicals, even at pH 12, are insufficient to destroy *C. albicans* unless supplemented by heat treatment (Figure 1–3). Moreover, it revealed that a buffer with SDS at RT, in the presence or absence of Proteinase K, effectively permeabilizes *C. albicans* cell wall but is unable to destroy the whole cell structure (Figure 4–5). Lastly, enzymatic reagents such as Zymolyase, on the other hand, provides an effective RT solution as it can fully destroy the cellular structure of *C. albicans* without supplemental heat, chemical, or physical treatment (Figure 6).

An interesting phenomenon in which nearly all *C. albicans* cells remain both AO- and PI-positive for a long period of time in the pH- and detergent-based studies was not observed in the Zymolyase experiment (Figures 1, 4, 6). Perhaps, the *C. albicans* cell structure is being destroyed quickly, and the biophysiological transition from healthy live cell status to permeabilized to loss of complete structure is not captured. A time-lapse study of live cells will be needed to address such a possibility. It is worth mentioning that some lysis buffers may not be compatible with direct mixing to the AO/PI dyes, and further process optimization (e.g., dilution or removal of lysis buffer) would be necessary to minimize potential chemical interference between the buffer and the dyes. It is also important to select appropriate marker(s) for quantification. For example, PI-marker cannot be used for dead cell counting in the Zymolyase study as cell structure was destroyed and those PI-positive clusters were likely the sticky genomic material left in the sample matrix. In summary, the Cellometer™ X2 fluorescent cell viability counter is feasible for performing quick and direct lysis efficiency studies.

## Acknowledgments

We thank Dr. Zhi-xiang Lu for his critical comments and invaluable revisions of this study.

